# Genome-wide survey of *Pseudomonas aeruginosa* PA14 reveals a role for the glyoxylate pathway and extracellular proteases in the limited utilization of mucin

**DOI:** 10.1101/091462

**Authors:** Jeffrey M. Flynn, Chi Phan, Ryan C. Hunter

## Abstract

Chronic airway infections by the opportunistic pathogen *Pseudomonas aeruginosa* are major cause of mortality in cystic fibrosis (CF) patients. While this bacterium has been extensively studied for its virulence mechanisms, biofilm growth and immune evasion within the CF airways, comparatively little is known about the nutrient sources that sustain its growth *in vivo*. Respiratory mucins represent a potentially abundant bioavailable nutrient source, though we have recently shown that canonical pathogens inefficiently use these host glycoproteins as a growth substrate. Yet, given that *P. aeruginosa*, particularly in its biofilm mode of growth is thought to grow slowly *in vivo*, the inefficient use of mucin glycoproteins may have relevance to its persistence within the CF airways. To this end, here we use whole genome fitness analysis combining transposon mutagenesis with high throughput sequencing (TnSeq) to identify genetic determinants required for *P. aeruginosa* growth using intact purified mucins as a sole carbon source. Our analysis reveals a biphasic growth phenotype, during which the glyoxylate pathway and amino acid biosynthetic machinery are required for mucin utilization. Secondary analyses confirmed the simultaneous liberation and consumption of acetate during mucin degradation and revealed a central role for the extracellular proteases LasB and AprA. Together, these studies identified the *P. aeruginosa* genes required for mucin-based nutrient acquisition and reveal a host-pathogen dynamic that may contribute to its persistence within the CF airways.

## INTRODUCTION

Cystic fibrosis (CF), a common and lethal autosomal recessive disease, results from mutations in the gene encoding the cystic fibrosis transmembrane conductance regulator (CFTR) protein (1). Impaired CFTR function leads to abnormal transepithelial ion transport and a thickened, dehydrated mucus layer that overlays the epithelium of several organs, including the lungs (2, 3). Airway mucus obstruction results in impaired mucociliary clearance, facilitating chronic colonization by a complex bacterial community. The ensuing inflammatory response leads to bronchiectasis, progressive lung damage, and eventual respiratory failure (2). Despite the recent surge in studies describing a polymicrobial etiology of CF, *P. aeruginosa* continues to be widely recognized as the primary driver of disease progression (4). This opportunistic pathogen can reach densities of 10^7^-10^9^ cfu gm-1 of sputum, particularly in late stages of disease, suggesting that the mucus environment of the lower airways provides them with an ideal growth environment (5, 6). A deeper understanding of this milieu, and its ability to sustain *P. aeruginosa* growth *in vivo* is critically important to improved disease management.

The mucosal layer covering the respiratory epithelium is a complex mixture of water, salts, protein, lipid, nucleic acids, mucins and lower molecular mass glycoproteins (7). The gel-forming mucins, MUC5B and MUC5AC, are highly glycosylated (70-80% by weight) polypeptides that form a cross-linked, hydrated gel and comprise the major macromolecular constituents of the mucus layer. These mucins protect the underlying airway mucosa as they provide a first line of innate immune defense against pathogens and environmental toxins (8). In CF, however, the hypersecretion and accumulation of mucins, their increased viscosity and impaired clearance compromises their role in host defense and are thought to contribute to disease pathophysiology (9-12). For example, *P. aeruginosa* has mucin-specific adhesins that mediate surface attachment by recognizing sialic acid and N-acetylglucosamine residues (13-15). Moreover, studies have shown that CF mucins are bound more efficiently by *P. aeruginosa* relative to non-CF mucins (15). Various reports have also described the role of mucins in stimulating bacterial surface motility, biofilm and aggregate formation, and the transition between biofilm and free-swimming states (16-20). In addition, the direct interaction of mucins with respiratory pathogens is thought to serve as a signaling event for the induction of various virulence factors (21).

Mucins also represent an abundant, host-derived nutrient reservoir for pathogens to utilize. In fact, all mucosal sites throughout the human body harbor both commensal and pathogenic organisms that possess mucin-degrading enzymes capable of deriving nutrients from host (22-24). Indeed, several studies have described the degradation and utilization of mucins by CF-associated microbiota (25-29). For example, multiple studies have shown that *P. aeruginosa* and other pathogens (e.g. *Burkholderia cepecia* complex, Bcc) harbor mucin sulfatase activity capable of utilizing sialylated mucin oligosaccharides as a source of sulfur (26, 28). Others have shown that Bcc isolates possess mucinase activity that can sustain pathogen growth using mucins as a sole carbon source (29). These studies emphasize the potential importance of mucins as a growth substrate for the initiation and maintenance of CF infections *in vivo*.

To investigate respiratory mucin-bacterial interactions in further detail, artificial sputum media (ASM) is commonly used to mimic the *in vivo* nutritional environment of the diseased airways. Among its constituents, porcine gastric mucin (PGM) is incorporated as a model carbon substrate because of its cost, ease of preparation, and similarity to human tracheal mucins in carbohydrate composition (30, 31). Use of PGM-based “sputum” media has generated insight into *P. aeruginosa* physiology under conditions relevant to the CF airways. For example, Sriramulu et al. (32) demonstrated that when PGMs are omitted from ASM, *P. aeruginosa* growth is limited, which implicated mucins as an important carbon source. Others have shown that PGM-based ASM media yield similar *P. aeruginosa* transcriptional profiles as the same isolates grown on re-constituted CF patient sputum (33). However, we have recently revealed that when commercial mucins are prepared such that low molecular weight metabolites are removed, *P. aeruginosa* growth is slow and significantly impaired. Growth was significantly improved (>10-fold) when co-cultured with mucin-degrading anaerobes (34). This observation suggests an inefficient use of intact mucin glycoproteins by the primary CF pathogen; however, this limited growth may have relevance to the slow *in vivo* growth rates of *P. aeruginosa* reported previously (35). Motivated by this possibility, here we further characterized the growth of *P. aeruginosa* PA14 on intact PGM. In addition, we used transposon insertion sequencing (TnSeq; 36) to identify the mechanisms by which *P. aeruginosa* degrades and consumes mucins when provided as a sole carbon source. Among essential 1 loci were genes associated with the glyoxylate pathway and amino acid biosynthesis. Further characterization and mutant analyses confirmed that both acetate and amino acids are consumed throughout a biphasic growth pattern. These results suggest that the liberation of these metabolites from airway mucins may have potential implications for *P. aeruginosa* growth and persistence within the CF airways.

## MATERIALS AND METHODS

### Bacterial strains, plasmids and culture conditions

A list of bacterial strains and plasmids is provided in Table 1. All strains were routinely cultured in Luria-Bertani (LB, Difco) broth and LB agar. Gentamicin sulfate (50 μg ml^−1^) and 2,6-diaminopimelic acid (DAP, 250 μM) were added where necessary. Mucin minimal medium (MMM) was prepared as described previously (34). Briefly, porcine gastric mucins (Type III, Sigma-Aldrich) were dissolved in water to 30 g L^−1^ and autoclaved. Mucins were then dialyzed against ultrapure water with a 13kDa MWCO dialysis membrane, and clarified by centrifugation at 100,000 x *g* for 30 min. MMM was made by adding the specified amount of mucins to a defined medium containing 60mM KH_2_PO_4_ (pH 7.4), 90mM NaCl, 1mM MgSO_4_, and a trace minerals mix described elsewhere (37). The final MMM was also filter-sterilized (0.45 μm). For growth with glucose or casamino acids (CasAA), the MMM described above was used without mucin but with added glucose or CasAA at the specified concentrations. Minimal glucose cultures were also supplemented with 60 mM NH_4_Cl.

**Table 1.**
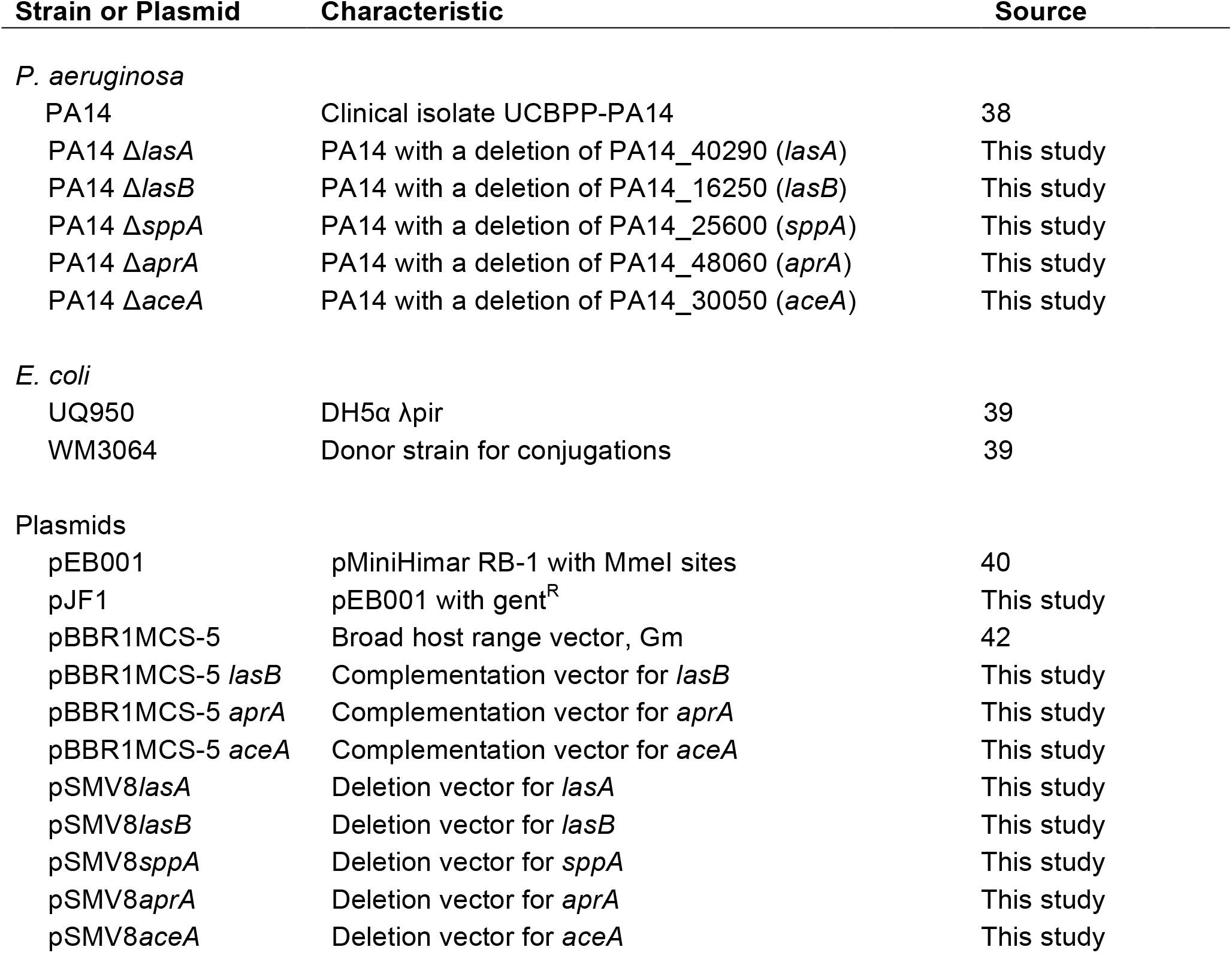
Bacterial strains and plasmids

| Strain or Plasmid | Characteristic | Source |
| --- | --- | --- |
| *P. aeruginosa* |  |  |
| PA14 | Clinical isolate UCBPP-PA14 | 38 |
| PA14 $\Delta lasA$ | PA14 with a deletion of PA14_40290 (*lasA*) | This study |
| PA14 $\Delta lasB$ | PA14 with a deletion of PA14_16250 (*lasB*) | This study |
| PA14 $\Delta sppA$ | PA14 with a deletion of PA14_25600 (*sppA*) | This study |
| PA14 $\Delta aprA$ | PA14 with a deletion of PA14_48060 (*aprA*) | This study |
| PA14 $\Delta aceA$ | PA14 with a deletion of PA14_30050 (*aceA*) | This study |
| *E. coli* |  |  |
| UQ950 | DH5α λpir | 39 |
| WM3064 | Donor strain for conjugations | 39 |
| Plasmids |  |  |
| pEB001 | pMiniHimar RB-1 with Mmel sites | 40 |
| pJF1 | pEB001 with gent<sup>R</sup> | This study |
| pBBR1MCS-5 | Broad host range vector, Gm | 42 |
| pBBR1MCS-5 *lasB* | Complementation vector for *lasB* | This study |
| pBBR1MCS-5 *aprA* | Complementation vector for *aprA* | This study |
| pBBR1MCS-5 *aceA* | Complementation vector for *aceA* | This study |
| pSMV8*lasA* | Deletion vector for *lasA* | This study |
| pSMV8*lasB* | Deletion vector for *lasB* | This study |
| pSMV8*sppA* | Deletion vector for *sppA* | This study |
| pSMV8*aprA* | Deletion vector for *aprA* | This study |
| pSMV8*aceA* | Deletion vector for *aceA* | This study |

### Growth on mucins

For growth assays, a colony was picked from freshly streaked LB agar, inoculated into LB broth, and allowed to grow for 16 h. Strains were washed twice with phosphate buffered saline before inoculation into 5 mL of MMM to an initial OD_600_ of ~0.05. Tubes were shaken continuously at 200 RPM at 37°C and OD_600_ was monitored over time using a Spectronic 20 spectrophotometer (Bausch and Lomb). 100 μL aliquots were removed throughout growth where specified, immediately frozen at −80°C and stored for downstream analysis.

### Protein and acetate quantification

Acetate was measured using an Acetate Colorimetric Assay Kit (Biovision) and protein content was measured using Qubit fluorometry (Thermo-Fisher) according to manufacturers protocols.

### TnSeq

TnSeq protocols were based on previously published methods (36, 40). First, the previously modified mariner transposon (41) was further modified to replace the kanamycin resistance cassette with a gentamicin cassette for use in *P. aeruginosa*. Briefly, the plasmid containing the transposon and transposase (pEB001)(40) was used as template in a PCR using TnSwap1 and TnSwap2 (a list of primers is shown in Table S1) to generate a fragment containing the whole pEB001 plasmid lacking a selectable marker. Next, the gene encoding gentamicin acetyltransferase (gent^R^) was amplified from pBBR1MCS-5 (42) using GentR_Fwd and GentR_Rev primers. Gibson assembly (43) was used to combine gent^R^ with the linearized pEB001 plasmid generated by PCR above to create pJF1. Next, pJF1 was transformed into the conjugative mating strain WM3064 (39) and a saturating transposon mutant library was generated by mating *P. aeruginosa* PA14 with WM3064 (containing pJF1) on LB + DAP agar plates followed by selection on LB gentamicin (50 μg mL^-1^). Mutants (~70,000) were pooled en masse in LB + 15% glycerol and were frozen in 100μL aliquots.

Outgrowth experiments are described in Figure 2. Briefly, one 100 μL aliquot was pelleted and frozen for gDNA extraction (“Parent”). A second glycerol stock was thawed, washed twice in PBS before inoculation into 5mL of MMM to an OD^600^ of 0.02. Cultures were shaken at 250 RPM at 37°C and OD^600^ was monitored until the cells reached ~0.4. 0.25mL of this culture was then used to inoculate a fresh 5mL of MMM and allowed to grow to ~0.4 (representing approximately 10 doublings, “1^st^ Phase”). A second culture was prepared by inoculating an additional 100μL glycerol stock into filter-sterilized spent 19 growth medium from the 1^st^ phase, followed by outgrowth to ~0.4 OD. As described above, 0.25mL was 20 used to inoculate a fresh 5mL of spent medium and allowed to grow again to ~0.4 (~10 doublings, “2^nd^ phase”). Bacterial cells were harvested by centrifugation and frozen at −80°C.

**Figure 1.**
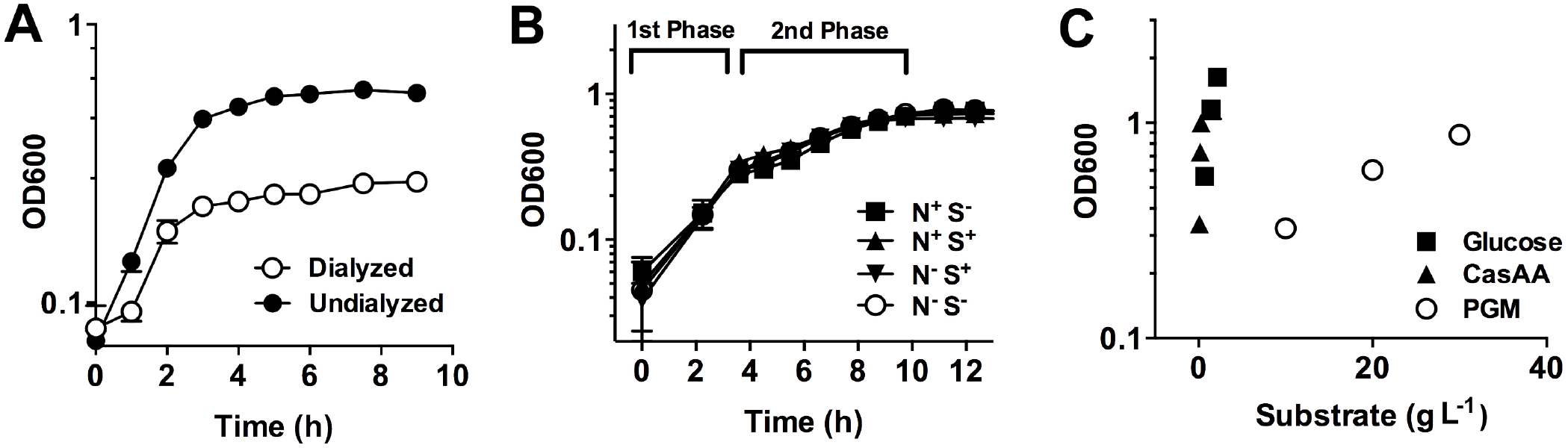
*P. aeruginosa* utilizes mucins inefficiently in a biphasic growth pattern. (A) PA14 growth on dialyzed PGM (10 g L-1) achieves half the final density relative to undialyzed preparations. (B) PA14 displays identical biphasic growth in the presence and absence of sulfur (MgSO4) and nitrogen (NH4Cl) 6 supplements when growing with 30 g L-1 PGM. (C) PA14 growth yields (g L-1 OD-1) on glucose and casamino acids far exceeded yield obtained with dialyzed PGM.

**Figure 2.**
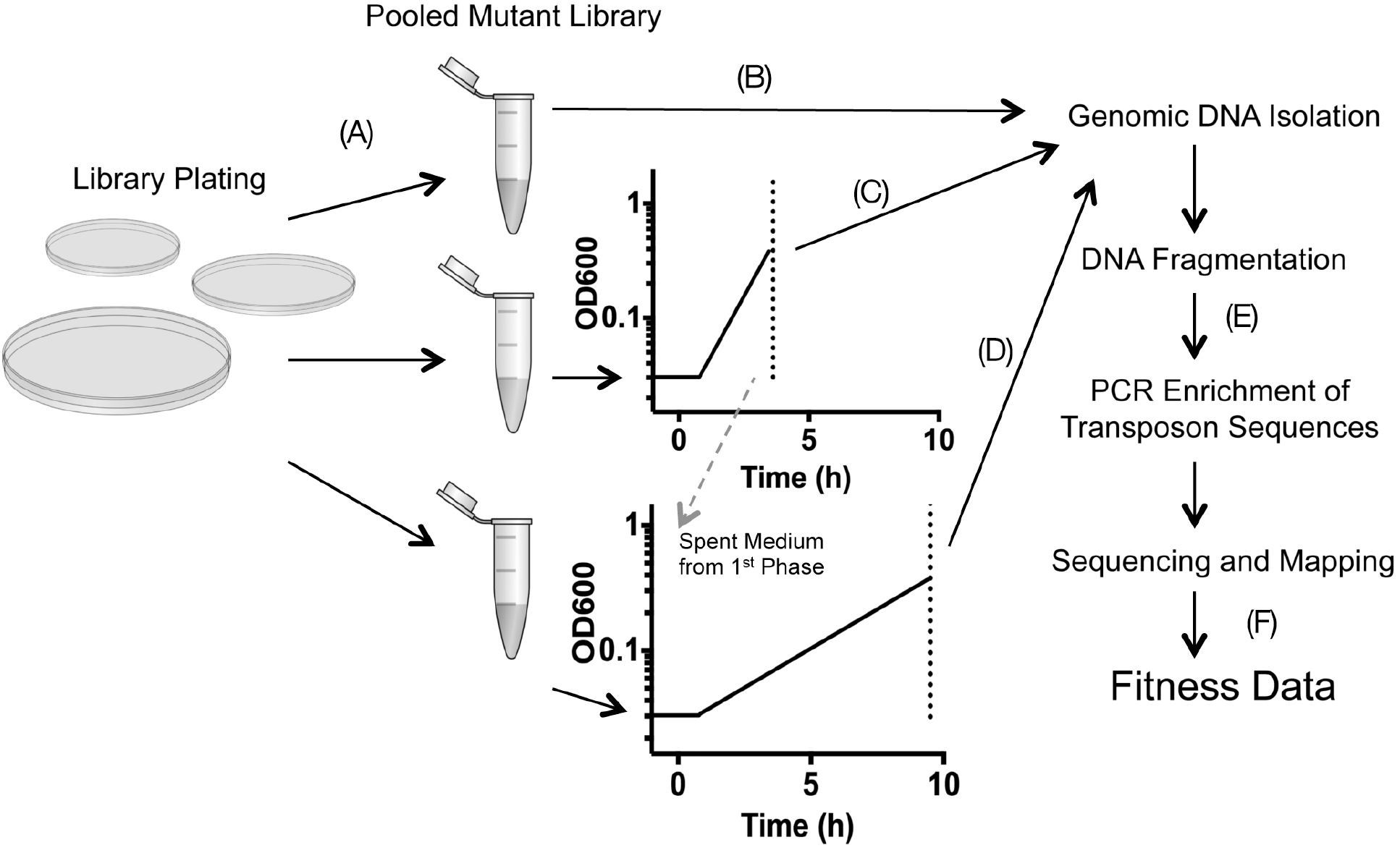
TnSeq experimental approach. (A) a transposon insertion library of PA14 is first constructed in which each mutant colony contains a single transposon insertion in its genome. (B) DNA is isolated from one aliquot of the pooled library, while two others are used as the source inoculum for two conditions under which selection is performed: (C) minimal mucin medium, and (D) filter sterilized mucin medium from (C) in which nutrients have been depleted. Genomic DNA is also isolated from each recovered culture. (E) Fragmented DNA is then PCR amplified generating bacterial-specific sequences flanked by Illumina-specific sequences with unique barcodes. (F) Sequence reads are then assigned to each selection conditions based on their barcode identifier, mapped to the PA14 genome counted, and used to calculate a fitness score of each transposon insertion.

Genomic DNA (gDNA) was extracted from frozen cell pellets using the Wizard Genomic DNA Purification Kit (Promega), cut with MmeI restriction enzyme (New England Biolabs) and treated with calf intestinal phosphatase (New England Biolabs). Oligonucleotide adapters (Table S1) with a 3’-NN overhang containing appropriate sequences for Illumina sequencing were ligated to the resulting fragments using T4 ligase (New England Biolabs). Adapters were barcoded with a unique 4 bp sequence 1 to enable multiplexing. PCR was then performed using ligation reactions as template with primers specific for the inverted repeat region on the transposon and the ligated adapter (P1_M6_MmeI and Gex_PCR_Primer, respectively). These primers introduce sequences suitable for direct sequencing in an Illumina flowcell. The resulting reaction product (120 bp) was purified by gel extraction using a PureLink Quick Gel Extraction Kit (LifeTechnologies) and sent to the University of Minnesota Biomedical Genomics Center for sequencing on the Illumina HiSeq 2500 platform (single end, 50 bp). Downstream analysis was performed on the Galaxy server (44-46) at the Minnesota Supercomputing Institute. Reads were split by unique barcode and trimmed to obtain the sequence adjacent to the transposon insertion. The resulting sequence was mapped to the PA14 genome (Genbank GCA_000014625.1) using Bowtie, discarding reads with greater than 1bp mismatch and sequences mapping to multiple locations. Insertions within the first 5% and last 10% of the open reading frame were also discarded to provide greater assurance that the insertion resulted in a null mutation. With these parameters, the Parent, 1^st^ 13 phase, and 2^nd^ phase had 27 million, 17 million, and 10 million total mapped hits, respectively. Genes with fewer than 10 unique insertions in the parent library were removed from analysis to limit their associated variability. Fitness values for each gene were then scored by computing a hit-normalized fold change for each growth condition (log^2^(outgrowth library hits / parent library hits)). This calculation identifies the fold-change in transposon mutant abundance between the parent and outgrowth populations (*i.e.* for a gene with a score of −2, there were 4X the number of cells containing a mutation in the parent library compared to the outgrowth). Negative fold-changes signified mutations with fitness defects under the outgrowth condition.

### Genetic manipulation

In-frame, markerless deletions in PA14 were generated using established homologous recombination techniques. Plasmids (Table 1) were derived from the suicide vector pSMV8 (39) and manipulated using standard molecular biology protocols with *E. coli* DH5α(UQ950). For deletion constructs, 1000 bp regions flanking the gene to be deleted (including 3-6 codons of the beginning and end of the genes) were amplified by PCR using primers listed in Table S1. Fragments were joined and cloned into pSMV8 digested with SpeI and XhoI using Gibson assembly and chemically transformed into UQ950. Positive ligations were screened by PCR, transformed into *E. coli* strain 1 WM3064 and mobilized into PA14 by conjugation. Recombinants of PA14 were selected for on LB agar plates containing gentamicin (Gm), and double recombinants were selected for on LB agar containing 6% sucrose. Complementation vectors were constructed by cloning the gene (*lasB, aprA,* and *aceA*) with an additional ~50 bp upstream of the start site followed by ligation into pBBR1MCS-5. Briefly, fragments were amplified by PCR using gene specific primers (Table S1). Resulting fragments were gel-purified and digested using the restriction enzymes KpnI, SacI, and XhoI where appropriate. After digestion, purified fragments were combined with digested pBBR1MCS-5, ligated using T4 ligase and transformed into UQ950. Positive transformants were screened by PCR and confirmed by sequencing. Complementation constructs were mated into PA14 via conjugation using WM3064 as described above. Successful matings were selected for by plating on LB agar containing gentamicin. All constructs and deletions were verified by sequencing.

## RESULTS

### PA14 exhibits an inefficient, biphasic growth phenotype using mucins as a growth substrate

To assess the ability of PA14 to break down and metabolize intact mucins, porcine gastric mucin (PGM) was prepared by autoclaving followed by dialysis to remove low molecular weight metabolites. This ensured that any observed growth was due to the use of large, intact glycoproteins. PA14 grew to a density of 0.6 in 10 g L^−1^ undialyzed mucins; however, when PGM was dialyzed, density fell to 0.25 (Fig 1A). These data suggest that over half of the bioavailable nutrients for PA14 in the undialyzed mucin preparation were composed of small metabolites.

To facilitate the study of *P. aeruginosa* growth on intact mucins, we then increased the PGM content in the culture medium to 30 g L^−1^, well above physiological concentrations. Under these 23 conditions, a biphasic growth pattern was observed. Growth up to 0.35 OD (“1^st^ Phase”) proceeded with 24 a doubling time approximately 2X that of growth from 0.35 to 0.8 (“2^nd^ phase”)(Fig. 1B), which reached its peak cell density after 11 hours. To determine if there was an unfulfilled nutrient requirement that limited *P. aeruginosa* growth in either growth phase, we then grew PA14 on PGM with and without supplements of sulfur (magnesium sulfate, 1mM) and nitrogen (ammonium chloride, 60mM)(Fig. 1B). Neither 1 supplement stimulated an increase in cell density, suggesting that the limiting nutrient of our minimal mucin medium (MMM) was likely carbon.

To assess the efficiency of PA14 to utilize PGM relative to other carbon sources (*e.g.*, the constituent amino acids and sugars in mucin glycoproteins), we then compared growth yields of PA14 when grown on glucose, casamino acids (CasAA), and PGM alone (Fig. 1C). On a gram-per-OD basis, PA14 obtained ~25X the density using glucose and ~100X the density using CasAA relative to dialyzed mucin (1.28, 0.3, and 34 g L^−1^ OD^−1^ for glucose, CasAA, and mucin, respectively). This limited growth yield of PA14 on mucins relative to other carbon sources underscores the inefficient use of intact mucin glycoproteins by *P. aeruginosa*.

### TnSeq reveals a need for glyoxylate bypass in mucin utilization

Despite the inefficient growth of *P. aeruginosa* on mucins, any breakdown and metabolism of these glycoproteins may be relevant to pathogen growth and persistence within the CF airways. Therefore, we sought to determine the mechanisms by which PA14 utilizes intact mucins in the absence of a complex, mucin-degrading bacterial community (34). To do so, we used a high-throughput transposon insertion sequencing approach, TnSeq, to identify the genetic requirements for *P. aeruginosa* throughout its biphasic growth phenotype described in Fig. 2B. This method allows for a comprehensive, single-culture mutant screen using a pooled transposon library, outgrowth of that library under a set of selective conditions, followed by Illumina sequencing to provide a semi-quantitative measure of fitness for each gene. Using the sequencing output, a fitness score for transposon insertions at each genetic locus can be calculated. We applied this technique to study PA14 growth under selection in the first (rapid) growth phase, and the second (slow) growth phase for ~10 generations each.

Our experimental approach is shown in Fig. 2. A library of ~70,000 transposon mutants were isolated on LB agar plates, pooled, and frozen. To perform the selection experiments, an aliquot was thawed and used to inoculate the MMM growth medium, and the remaining culture was then pelleted and frozen for genomic DNA isolation. Two medium conditions were then tested for selection: (i) dialyzed minimal mucin medium (MMM) and (ii) MMM depleted of the “1^st^ phase” carbon source. To achieve the 1 latter, PA14 was grown in MMM until an OD^600^ of 0.4 and the spent medium was filter sterilized. Media were then inoculated to 0.02 OD from the parent library and were allowed to grow to OD^600^ ~0.4. The resulting cultures were then re-inoculated to 0.02 OD and allowed to grow again to 0.4. Total growth was equal to approximately 10 doublings. Genomic DNA directly adjacent to each transposon insertion was then sequenced and insertion sites for each genetic locus in both the outgrowth and parent libraries were quantified. Using the relative abundance of transposon insertions at each locus, fitness scores were calculated for the essentiality of each gene in the two outgrowth conditions. The parent library contained 59,380 unique transposon insertion sites with an average of 8.7 insertions in the central 85% of the coding region. To calculate fitness values for each gene under a given selection condition, the number of reads mapped to a given gene in the outgrowth library were divided by the number of reads in the parent library, followed by a base-two logarithm conversion to calculate fold-changes between populations. After this transformation, a score of −2 or less was interpreted as a significant fitness defect.

The data revealed that multiple genes required for the first growth phase were not required for the second growth phase (Table 2, Table S2). By contrast, no genes were identified that were required for the second growth phase but not required for the first. In the first growth phase, multiple transposon insertions in genes encoding glyoxylate pathway enzymes had decreased fitness. These included *aceA,* encoding isocitrate lyase, and *glcB,* encoding malate synthase. Fitness defects in these transposon mutants suggested that the primary carbon source during rapid growth (1^st^ phase) was either a 2-carbon compound (*e.g.* acetate) or fatty acids, both of which require the action of the glyoxylate pathway for catabolism and growth. In addition to glyoxlyate requirements, PA14 also required multiple genes encoding amino acid biosynthesis enzymes (*arg*, *cys*, *ilv*, *leu*, *met*, *trp*). This requirement suggested that despite being grown on a carbon-rich glycoprotein, any liberation of amino acids from the mucin polypeptide was not rapid enough to satisfy the nutritional requirements of *P. aeruginosa* during the first growth phase.

**Table 2.**
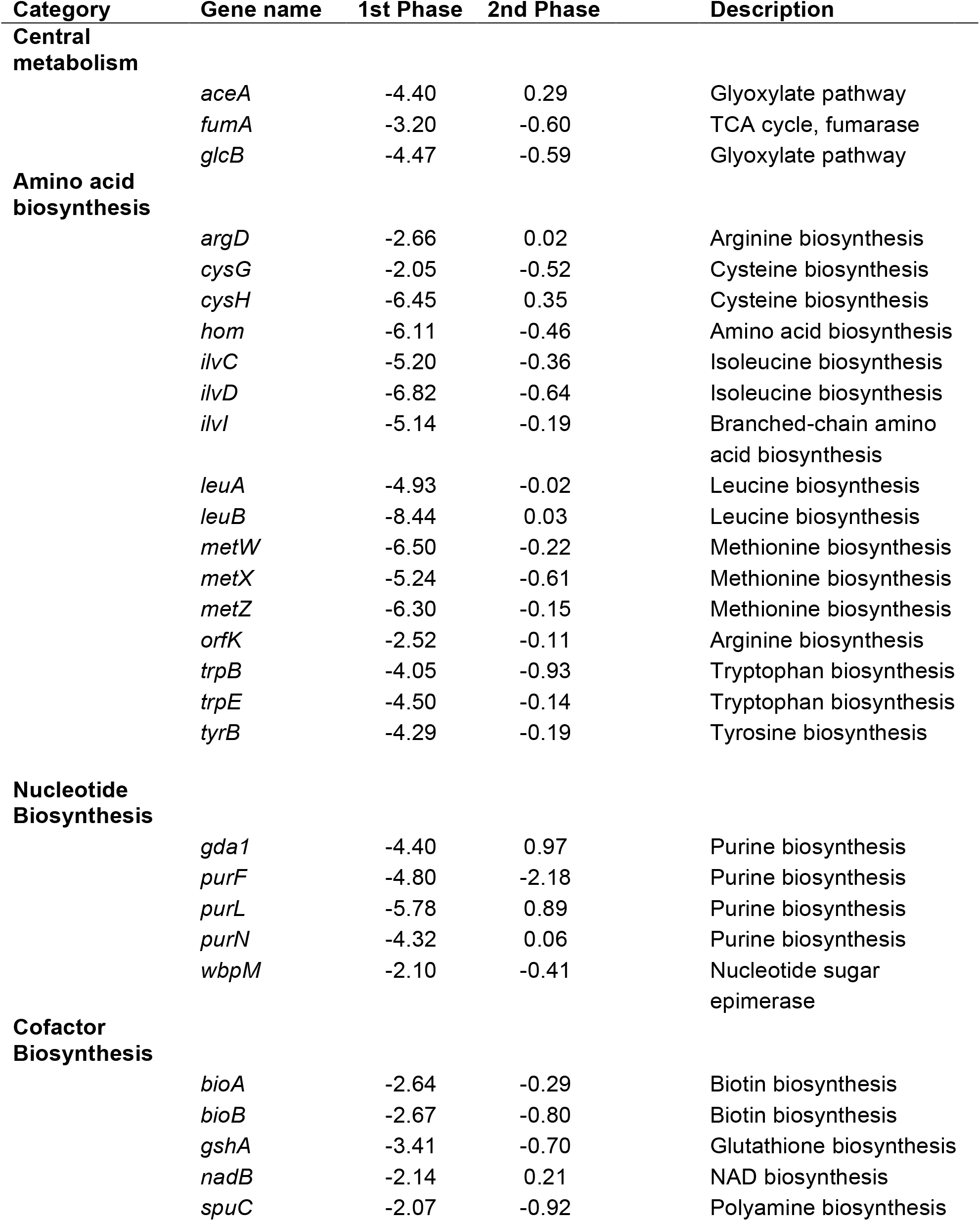
TnSeq fitness values.

| Category | Gene name | 1st Phase | 2nd Phase | Description |
| --- | --- | --- | --- | --- |
| **Central metabolism** | *aceA* | -4.40 | 0.29 | Glyoxylate pathway |
|  | *fumA* | -3.20 | -0.60 | TCA cycle, fumarase |
|  | *glcB* | -4.47 | -0.59 | Glyoxylate pathway |
| **Amino acid biosynthesis** | *argD* | -2.66 | 0.02 | Arginine biosynthesis |
|  | *cysG* | -2.05 | -0.52 | Cysteine biosynthesis |
|  | *cysH* | -6.45 | 0.35 | Cysteine biosynthesis |
|  | *hom* | -6.11 | -0.46 | Amino acid biosynthesis |
|  | *ilvC* | -5.20 | -0.36 | Isoleucine biosynthesis |
|  | *ilvD* | -6.82 | -0.64 | Isoleucine biosynthesis |
|  | *ilvI* | -5.14 | -0.19 | Branched-chain amino acid biosynthesis |
|  | *leuA* | -4.93 | -0.02 | Leucine biosynthesis |
|  | *leuB* | -8.44 | 0.03 | Leucine biosynthesis |
|  | *metW* | -6.50 | -0.22 | Methionine biosynthesis |
|  | *metX* | -5.24 | -0.61 | Methionine biosynthesis |
|  | *metZ* | -6.30 | -0.15 | Methionine biosynthesis |
|  | *orfK* | -2.52 | -0.11 | Arginine biosynthesis |
|  | *trpB* | -4.05 | -0.93 | Tryptophan biosynthesis |
|  | *trpE* | -4.50 | -0.14 | Tryptophan biosynthesis |
|  | *tyrB* | -4.29 | -0.19 | Tyrosine biosynthesis |
| **Nucleotide Biosynthesis** | *gda1* | -4.40 | 0.97 | Purine biosynthesis |
|  | *purF* | -4.80 | -2.18 | Purine biosynthesis |
|  | *purL* | -5.78 | 0.89 | Purine biosynthesis |
|  | *purN* | -4.32 | 0.06 | Purine biosynthesis |
|  | *wbpM* | -2.10 | -0.41 | Nucleotide sugar epimerase |
| **Cofactor Biosynthesis** | *bioA* | -2.64 | -0.29 | Biotin biosynthesis |
|  | *bioB* | -2.67 | -0.80 | Biotin biosynthesis |
|  | *gshA* | -3.41 | -0.70 | Glutathione biosynthesis |
|  | *nadB* | -2.14 | 0.21 | NAD biosynthesis |
|  | *spuC* | -2.07 | -0.92 | Polyamine biosynthesis |

The second, slower growth phase showed no defects conferred by single transposon insertions when compared to the first growth phase alone. These results, in sharp contrast to the first phase, are indicative of a diverse nutritional substrate pool whereby the disruption of a single gene does not confer 1 a growth defect. For example, PA14 did not require any specific amino acid biosynthetic genes, suggesting that a complete mix of amino acids were liberated from mucins and were available as ‘community goods’. Under these conditions, transposon insertions that would otherwise result in a growth-inhibited phenotype for an auxotophic mutant in pure culture might not confer a defect in a community of pooled transposon mutants with mixed abilities. Given the lack of amino acid biosynthetic genes required for the second growth phase, we hypothesized that amino acids were being liberated from mucins and serving as the primary source of carbon and energy.

### PA14 liberates and consumes acetate in the first phase of growth and consumes amino acids throughout growth

Given the requirement for the glyoxylate pathway, we hypothesized that acetate may be liberated from mucin. To assess this, we monitored acetate concentrations in the medium throughout growth. Though no acetate was initially present in the growth medium, it rapidly accumulated in the culture supernatant during the first growth phase (Fig. 3A). Its concentration increased until the transition between the first and second growth phase (~0.3 OD), where it was rapidly consumed. This result is consistent with the glyoxylate pathway being required for the first growth phase in which 2-carbon compounds serve as a primary carbon source.

**Figure 3.**
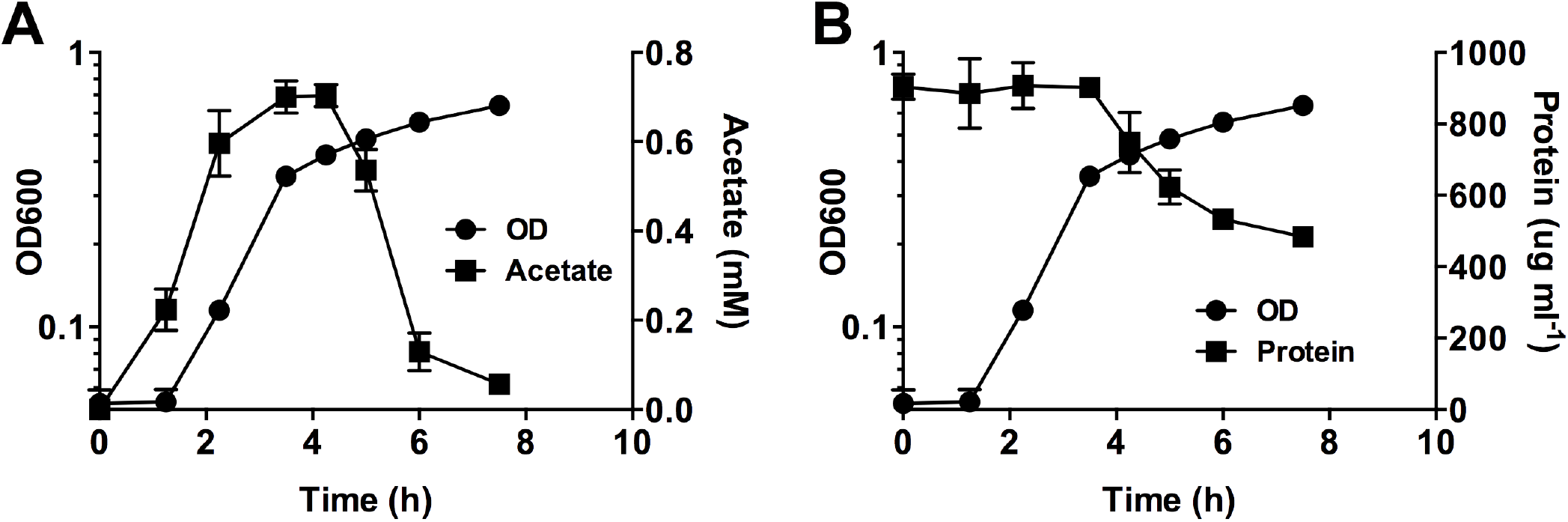
Acetate and protein content during mucin growth. (A) Acetate accumulates in the growth medium followed by its rapid consumption by *P. aeruginosa*. (B) Total protein also decreases in the second growth phase, suggesting amino acid liberation from the mucin polypeptide.

Given that no amino acid biosynthetic genes were essential in the second growth phase, we predicted that amino acids were liberated via degradation of the mucin glycoprotein. We therefore quantified protein content (excluding free amino acids) in the medium supernatant (Figure 3B). Interestingly, protein concentration remained unchanged in the first phase of growth, and rapidly declined in the second phase. Taken together, these observations demonstrate a sequential use of carbon sources (acetate, followed by amino acids) derived from the mucin glycoprotein.

### Growth of *aceA*, *lasB* and *aprA* mutants demonstrate growth defects in PGM

To confirm the importance of acetate and the glyoxylate pathway in the first phase of growth, a non-polar, markerless deletion was made in *aceA,* encoding isocitrate lyase, which catalyzes the first step in the glyoxylate pathway. Any strain lacking this committed step would not be able to use acetate as a sole carbon 1 source because it would be unable to correctly balance carbon requirements in the TCA cycle. Consistent with this limitation, the *aceA* mutant demonstrated a marked growth defect relative to WT and reached a lower final density (Fig. 4A, Fig. S1). The incomplete abolishment of growth suggests that the Δ*aceA* mutant is capable of obtaining carbon via other means but that the glyoxylate pathway defect restricts its growth and overall yield.

**Figure 4.**
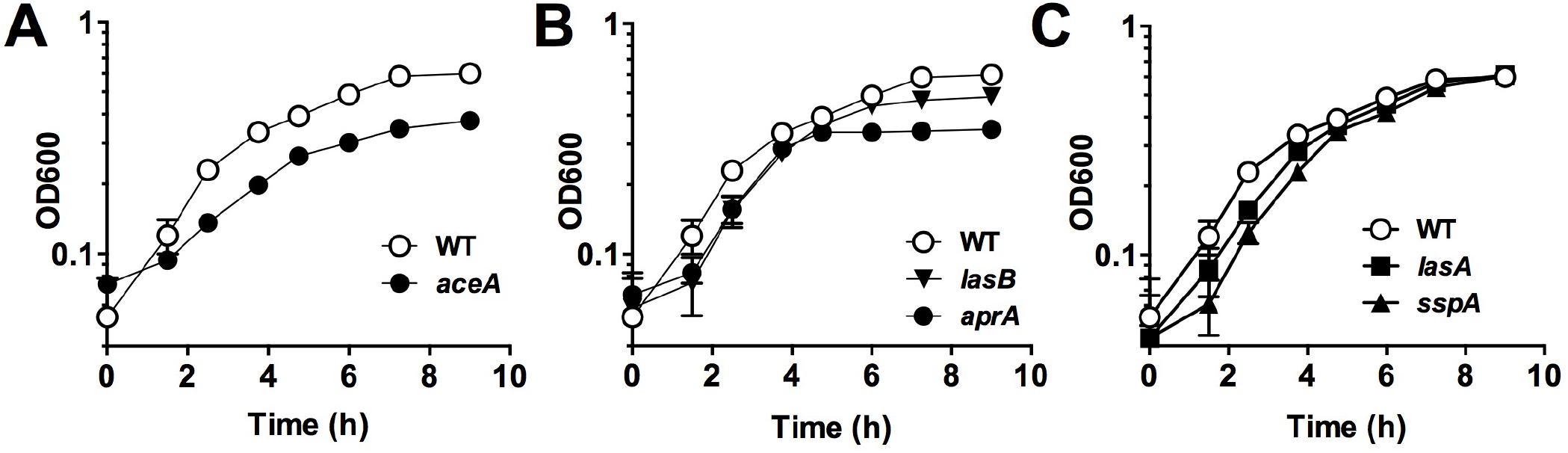
AceA, LasB and AprA are used during mucin growth. (A) Deletion of isocitrate lyase (AceA) confers a partial growth defect in *P. aeruginosa* when provided mucins as the sole carbon source. The extracellular proteases (B) LasB and AprA, but not (C) LasA and SspA are used in the degradation and consumption of mucin polypeptides.

Protein consumption in the second growth phase suggested that one or more extracellular proteases were being employed to degrade the mucin polypeptide backbone. To address this possibility, we tested four extracellular proteases (LasA, LasB, AprA and SppA) that may be important for the degradation of extracellular peptides. Non-polar, markerless deletions were made in genes encoding each of these proteases and their growth was assayed versus WT in MMM (Fig. 4B,C). Interestingly, only mutants lacking LasB and AprA exhibited defects in the second phase of growth and lower final densities, while complementation with *lasB* and *aprA in trans* was able to restore the WT phenotype (Fig. S1). These data demonstrate that elastase (LasB) and alkaline protease (AprA) are used by *P. aeruginosa* during slow growth on mucins to utilize the polypeptide backbone as a carbon source.

## DISCUSSION

In contrast to *Streptococcus, Akkermansia, Bacteroides* and other bacterial genera that have an extensive repertoire of enzymes devoted to catabolizing host-associated glycoproteins in the gut and oral cavity (24, 47-50), the primary cystic fibrosis pathogen, *P. aeruginosa,* is not known to encode any glycosidases used in the breakdown of respiratory mucins. Indeed, here we demonstrate that strain PA14 uses intact mucins inefficiently compared to its carbohydrate and amino acid monomeric constituents (glucose and amino acids). However, when purified and dialyzed mucins were provided at high concentrations, we observed a slow, biphasic growth of PA14 to an appreciable density. This observation suggests that a even a partial breakdown of mucin glycoproteins may support the slow growth (35) and persistence of *P. aeruginosa* in the CF airways.

The limited degradation of porcine-derived mucins by strain PA14 was not unexpected; in the oral cavity, for example, diverse consortia of bacteria are required to completely breakdown salivary mucins 1 (50-51). Yet, our data are in contrast to previous studies that have implicated *P. aeruginosa* and another CF pathogen, *Burkholderia cenocepecia*, in the degradation of mucins within the airways (25, 29). We propose that this discrepancy may be explained by the limitations of mucin detection methods. Immunoblotting, a commonly used approach for evaluating mucin content, relies on anti-MUC5AC and anti-MUC5B antibodies directed towards their epitopes on the terminal, non-glycosylated regions (*i.e.* apomucin) of the macromolecule, as previously shown (52). Given that extracellular proteases (and not glycosidases) were found here to be essential for PA14 growth on PGM, we suspect that proteolytic degradation removes only the terminal polypeptide regions, leaving the bulk of the mucin glycoprotein intact. This limited degradation would give the impression of more extensive mucin breakdown via Western immunoblotting (52). Consequently, we favor the interpretation that mucinase activity previously ascribed to *P. aeruginosa* is likely due to terminal polypeptide degradation mediated by the extracellular proteases (LasB, AprA) described here (Fig. 4).

TnSeq analysis implicated a critical role for the glyoxylate shunt (*aceA, glcB*) in the generation and consumption of nutrients during the first phase of mucin growth. The requirement for this pathway implies a shift to C2 carbon metabolism, and led to the discovery of the use of acetate by *P. aeruginosa* during growth on PGM. The requirement for the glyoxylate pathway stems from the fact that carbon sources entering central metabolism below pyruvate must go through the TCA cycle to malate to be fed to gluconeogenic pathways. To correctly balance TCA cycle metabolites, the glyoxlyate pathway is required for the regeneration of four carbon intermediates (*e.g.* malate). While the glyoxylate shunt is also essential for fatty acid and lipid metabolism under nutrient-limited growth conditions, direct measurements of the culture supernatant confirmed that acetate accumulates in the growth medium followed by its consumption. We hypothesize that acetate is likely derived via deacetylation of sugars (N-acetylglucosamine, N-acetylgalactosamine and sialic acids) that decorate the polypeptide backbone., Further studies are required to determine the source and enzymes responsible for the accumulation of this metabolite.

Interestingly, the glyoxylate pathway has been implicated in various *in vivo* infection models. For example, an isocitrate lyase mutant (Δ*aceA*) of *P. aeruginosa* demonstrated significantly less virulence 1 and host tissue damage in a rat lung infection model (53). In addition, a Δ*aceA*Δ*glcB* double mutant of *P. aeruginosa* was cleared by 48h post-infection in a murine acute pneumonia model (54), yet showed no defective phenotypes in septicemia, underscoring its importance in the nutrient-limited environment of the lower airways. Son et al. (55) demonstrated that genes required for the glyoxylate cycle in *P. aeruginosa* are highly expressed within sputum derived from CF patients, while others have reported that *aceA* is more highly expressed in CF *P. aeruginosa* isolates compared to those derived from non-CF sources when grown *in vitro* (56). Notably, acetate has also been found at elevated concentrations within CF sputum and bronchoalveolar lavage fluid (34,57,58). Our data, when considered in the context of the aforementioned studies, support a role for the glyoxylate shunt in respiratory mucin degradation by *P. aeruginosa*. Moreover, since isocitrate lyase (AceA) and malate synthase (GlcB) have no known ortholog in humans, our data also provide further motivation to explore the glyoxylate shunt as a target for anti-Pseudomonal therapy, as previously suggested (59-62).

Though TnSeq did not identify extracellular proteases, mutant analysis demonstrated that *lasB* and *aprA*, encoding elastase B and alkaline protease, respectively, were also key enzymes required for mucin-based growth. Elastase B is a quorum sensing-regulated, prototypical virulence factor of *P. aeruginosa* (63, 64) and has been shown to degrade a wide variety of host proteins such as collagen, elastin, immunoglobulins and complement (65). Similarly, AprA (aeruginolysin) is a metallo-endopeptidase that has also been shown to degrade physiological substrates *in vivo* (65). While it is possible that other, yet-to-be identified proteases also contribute to mucin degradation, both LasB and AprA have been detected in abundance in the CF airways (66-69). Importantly, both *lasB* and *aprA* mutants have also shown attenuated virulence in animal infection models (70, 71), suggesting an important role for these proteases in lung disease pathogenesis. Given that both enzymes have also been implicated in bacterial keratitis, where colonization of the corneal layer was restricted in both LasB and AprA*-*deficient mutants (72, 73), we speculate that these two proteases are also important for *P. aeruginosa* nutrient acquisition and persistence at other mucin-rich sites of infection.

Understanding how respiratory pathogens adapt to the *in vivo* nutritional environment has important implications for the treatment of CF lung disease. As a step in this direction, this genome-wide 1 survey of mucin utilization strategies has provided a window into a potential mechanism of *P. aeruginosa* growth within the lower airways. While growth on mucins was limited, our data demonstrate that mucin degradation alone can support low densities of PA14, which may allow for bacterial persistence when nutrients are scarce (*e.g.* early stages of disease). We concede that the portrait of *in vivo* pathogen growth is never simple in a disease with an array of etiologies. It is likely that as airway infections evolve, the bacterial community and the host airway milieu change over time and between patients such that bioavailable nutrient pools are altered. Going forward, it will be important to consider how bacterial carbon acquisition strategies vary with disease states, and how they can be manipulated as a means of mitigating chronic CF airway infections.

## ACKNOWLEDGEMENTS

We acknowledge Caleb Levar and Chi-Ho Chan (Biotechnology Institute) for their assistance with TnSeq, and the Minnesota Supercomputing Institute (MSI). This work was supported by a Pathway to Independence Award from the National Heart Lung and Blood Institute to RCH (R00HL114862) and the National Center for Advancing Translational Sciences (UL1TR000114). JMF was support by a National Institutes of Health Lung Sciences T32 fellowship (#2T32HL007741-21) awarded through the National Heart Lung & Blood Institute and a Cystic Fibrosis Foundation Postdoctoral Fellowship (FLYNN16F0).

